# Pocket Restraints Guided by B-Cell Epitope Prediction Improves Chai-1 Antibody-Antigen Structure Modeling

**DOI:** 10.1101/2025.09.17.676770

**Authors:** Joakim Clifford, Morten Nielsen

## Abstract

The accurate prediction of antibody-antigen (AbAg) complexes is a key challenge for computational immunology, with applications in therapeutic antibody design and diagnostics. Current deep learning methods, such as AlphaFold and Chai, have the potential to generate high-confidence AbAg structures. However, these methods often fail to predict the correct AbAg structure, placing the antibody incorrectly on the antigen and converging on repeatedly predicting the same redundant binding mode. Here, we present **BepiPocket** and **DiscoPocket**, two simple approaches that integrate B-cell epitope prediction tools to guide antibody-epitope restraints during Chai-1 structure prediction. On a dataset of 1628 antibody-antigen complexes, we demonstrate that using the sequence-based predictor BepiPred-3.0 (BepiPocket) and the structure-based predictor DiscoTope-3.0 (DiscoPocket), we substantially improved both the accuracy and diversity of predicted AbAg complexes compared to standard Chai-1 modeling with random seed variation. We also find that a key driver in these performance gains is the antigen modeling accuracy. The software for both BepiPocket and DiscoPocket algorithms is freely available at https://github.com/mnielLab/BepiPocke

## Introduction

B cells are key components of the adaptive immune system, often providing long-term protection against pathogens and cancer. They are activated when B-cell receptors (BCRs)—membrane-bound antibodies—bind to specific antigens. The contact regions on the antigen and antibody are referred to as the epitope and paratope, respectively. Identifying B-cell epitopes is important for vaccine design (1), diagnostics (2), and therapeutic antibody development (3).

To reduce the time and cost of experimental screening, in silico epitope prediction methods are widely used to prioritize candidate epitopes. Most existing tools are antibody-agnostic, i.e. aim to identify antigen surface residues likely to engage with antibodies in general. Examples include BepiPred (4–6), DiscoTope (7, 8), and SEPPA (9). Although trained on epitope data annotated from antibody-antigen complexes, these models only integrate antigen sequence or structure information and do not explicitly incorporate antibody information. Even though the performance of these tools has improved over the last years, benchmarking these tools is inherently difficult. This is because only a few antigens, such as lysozyme or the SARS-CoV spike protein, have been studied extensively, with many solved antibody–antigen (AbAg) complexes. Most antigens are much less characterized, and available structures often capture antibodies bound to only a single region. This makes it challenging to evaluate antibody-agnostic tools, as predicted epitopes outside this known binding region may either be incorrect or simply lack experimental validation. Also, for many real life applications, one is more interested in identifying the binding target and binding location of a given antibody compared to identifying the likely B cell epitopes on a given antigen. Given this, a more precise and often more interesting task is therefore antibody-specific epitope prediction, which incorporates both antibody and antigen inputs to model this specificity (10). These methods have applications in autoimmune disease, cancer immunotherapy, and drug hypersensitivity, where identifying the exact target of an antibody can inform diagnosis or treatment.

Experimental techniques such as X-ray crystallography and cryo-electron microscopy (cryo-EM) can resolve AbAg structures at high resolution but are costly, time-consuming, and limited by practical constraints like crystallization (11, 12). Faster approaches such as phage display lack atomic detail (13). These challenges highlight the value of computational tools that can accurately predict antibody-specific epitopes. However, creating such tools is challenging due to the flexible and diverse nature of the antibody hypervariable complementarity-determining regions (CDRs) loops that mediate antigen binding. Early work by Culang et al. (14) and Jespersen et al. (15) aimed to address this using machine learning to predict antibody-antigen (AbAg) interactions based on sequential, physicochemical, and geometric features, using both antibody and antigen inputs. More recently, innovations in protein structure prediction, such as AlphaFold (16), offer a novel approach enabling the direct generation of full 3D AbAg complex structures from sequence. Our previous study showed that the structural accuracy of predicted AbAg with AlphaFold-2.3 is still lacking, but that the generated structural models are sensitive to the specific antibody-antigen pairing (17). That is, predictions made with incorrect antibody-antigen pairs rarely achieve the structural accuracy observed with the correct pair. This suggests that improving the structural accuracy of these tools could also improve the predictability for antibody specificity. Several new tools, including AlphaFold-3 (18), Chai (19), and Boltz (20), have recently been developed to improve the accuracy of structural predictions. While their antibody specificity has not been comprehensively evaluated, it is evident from these publications that the structural accuracy for AbAg complex predictions has improved compared to earlier models. A new feature introduced in recent tools such as Chai and Boltz is support for restraint-based structure prediction. Their publications have shown that this feature substantially improves the accuracy of predicted AbAg interfaces. However, these restraints typically require experimental data, which are often unavailable or costly to obtain. In this study, we demonstrate that in the absence of experimental data, antibody-agnostic B-cell epitope prediction tools—specifically BepiPred-3.0 and DiscoTope-3.0—can be used to define informative antibody-epitope restraints for use in Chai-1, offering substantial increases in the structural accuracy of predicted AbAg complexes.

## Results

### Solved structure pocket restraints substantially improve Chai-1 antibody–antigen structure prediction

A novel feature in recent structure prediction tools, such as Chai-1, is the support of restraint-based structure prediction, where spatial restraints—termed pocket restraints—can be applied to guide the modeling process. More specifically, a pocket restraint is a soft restraint (i.e., not strictly enforced) that any residue in one chain be in contact with a specific residue in another.

In this section, we investigated how such restraint-based structure prediction can be used to create antibody-epitope pocket restraints that improve the accuracy of AbAg structure prediction. As a baseline, we used Chai-1 without restraints or multiple sequence alignment (MSA) input— relying only on ESM (21) embeddings and a single seed, resulting in 5 predicted structures for each query AbAg. We applied this pipeline to sequences from a data set of 1,628 AbAg complexes. This dataset is a subset of AbAgs structures from our previous study Clifford et. al. (17), consisting of solved AbAg complexes deposited to the Protein Data Bank (PDB) biological assembly FTP server before February 2, 2023 (22). For details on the dataset and structure predictions refer to Methods. We then repeated predictions with pocket restraints derived from the corresponding solved structures, requiring the heavy chain or both chains to be within 10 Å of: (i) the top epitope residue with most antibody contacts, (ii) the top four residues, or (iii) all epitope residues. Structural confidence was measured using the default Chai-1 confidence metrics (0.8ipTM + 0.2pTM). We compute two success rates, defined as the percentage of AbAgs where the highest-confidence structure achieves a DockQ (23) score above a CAPRI threshold, and where any of the five predicted structures (Oracle) achieves a score above the same thresholds. CAPRI DockQ thresholds are defined as: acceptable (≥ 0.23), medium (≥ 0.49), and high (≥ 0.80).

The results of this analysis demonstrate that across all DockQ thresholds, including both antibody chains (red) in the restraint setting consistently improved performance over heavy chain-only (grey) and no-restraint (blue). Moreover, as expected, restraining more epitope residues generally yielded higher structural accuracy (darker red compared to lighter red) (**Figure 1A**).

**Figure 1.**
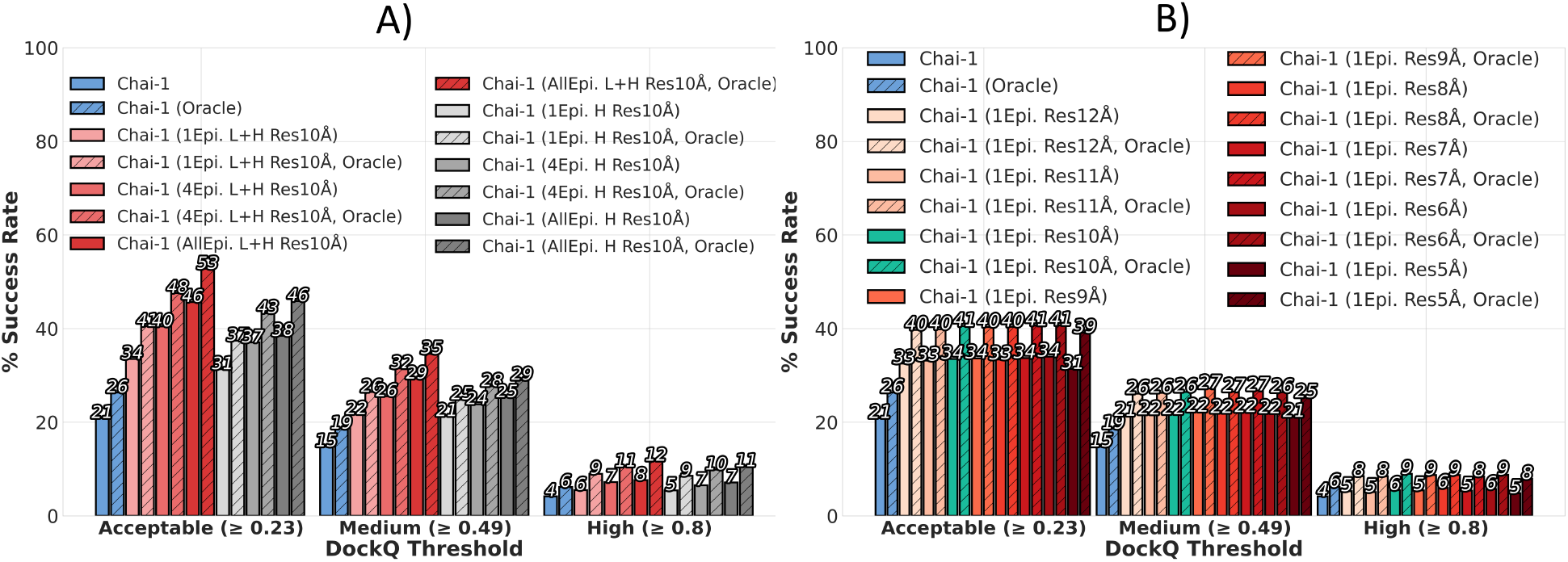
Investigation on how pocket restraint configurations, based on epitope contact information from solved structures, improves Chai-1’s structure prediction of 1,628 antibody–antigen (AbAg) complexes. **A)** Bar plot showing the percentage of AbAg predictions surpassing the CAPRI-standard DockQ accuracy thresholds— acceptable (≥ 0.23), medium (≥ 0.49), and high (≥ 0.80)—referred to as the success rate (y-axis). Results are shown for Chai-1 without pocket restraints (blue), with restraints requiring only the antibody heavy chain (grey), or both the light and heavy chains (red) to be within 10 Å of: (i) the epitope residue with the most antibody contacts in the corresponding solved structure, (ii) the top four such residues, or (iii) all epitope residues. Each method creates 5 structures per AbAg, and we show success rate when choosing the highest-confidence structure and the best-by-DockQ structure (Oracle). **B)** Bar plot showing DockQ success rates for Chai-1 predictions using pocket restraints that require both the light and heavy chains to be within 5, 6, 7, 8, 9, 10, 11, or 12 Å of the single epitope residue with the most antibody contacts.

We next evaluated how varying the restraint distance (5–12 Å) affected performance. A 5 Å cutoff consistently performed worst, while 12 Å performed second worst. The best results occurred between 6–11 Å, but with minimal variation within that range. For consistency and simplicity in downstream analyses, we selected a single epitope restraint distance of 10 Å for both light and heavy chain (teal) (**Figure 1B**).

### BepiPred-3.0 pocket restraints improve Chai-1 antibody-antigen structure prediction

In the previous section, we demonstrated that antibody-epitope pocket restraints, in the case of knowing the top epitope residue (the one with the most antibody contacts in the solved AbAg structure) can improve Chai-1 structure prediction of AbAg complexes. However, in most cases such structural epitope information is not available. We therefore next investigated if BepiPred-3.0—an antibody-agnostic B-cell epitope predictor—can identify informative restraint residues that substantially boost performance. We refer to this method as BepiPocket.

For the analyses in this section, we predicted structures for the 1,628 AbAgs in our dataset with Chai-1 using either 30 different seeds or 30 iterations using the BepiPocket algorithm to identify the pocket restraints. Both methods produce 30x5=150 structures per AbAg. Furthermore, in order to compute the antigen RMSD, we predicted 5 unbound (no antibody) antigen structures with Chai-1 using a single seed.

In short, the BepiPocket algorithm takes a list of antigen residues scored by BepiPred-3.0 (reflecting epitope likelihood) and iteratively selects the top-ranked residue in each Chai-1 run as the pocket restraint (for details, see Methods). Before using BepiPred-3.0 to guide structure prediction, we conducted a preliminary analysis comparing it to random scoring and two structure-based alternatives: DSSP relative surface accessibility (RSA) (24–26), and DiscoTope-3.0 (8). This analysis confirmed that BepiPred-3.0 ranked true epitope contacts from solved structures higher than the alternative methods, more reliably prioritizing residues involved in antibody binding. We estimated that after 30 runs, the true top epitope contact (1Epi; see Figure 1) would be selected in approximately 50% of AbAg predictions. A detailed description of this preliminary analysis is provided in Supplementary S1.

Based on this, we ran the Chai-1 for 30 runs per AbAg with BepiPred-3.0, calling the method BepiPocket. When compared to Chai-1 predictions run with 30 different seeds, BepiPocket demonstrated an improved performance, with 4% more AbAgs achieving acceptable or medium DockQ (CAPRI thresholds), and 2% more achieving high DockQ accuracy (**Figure 2A**).

**Figure 2.**
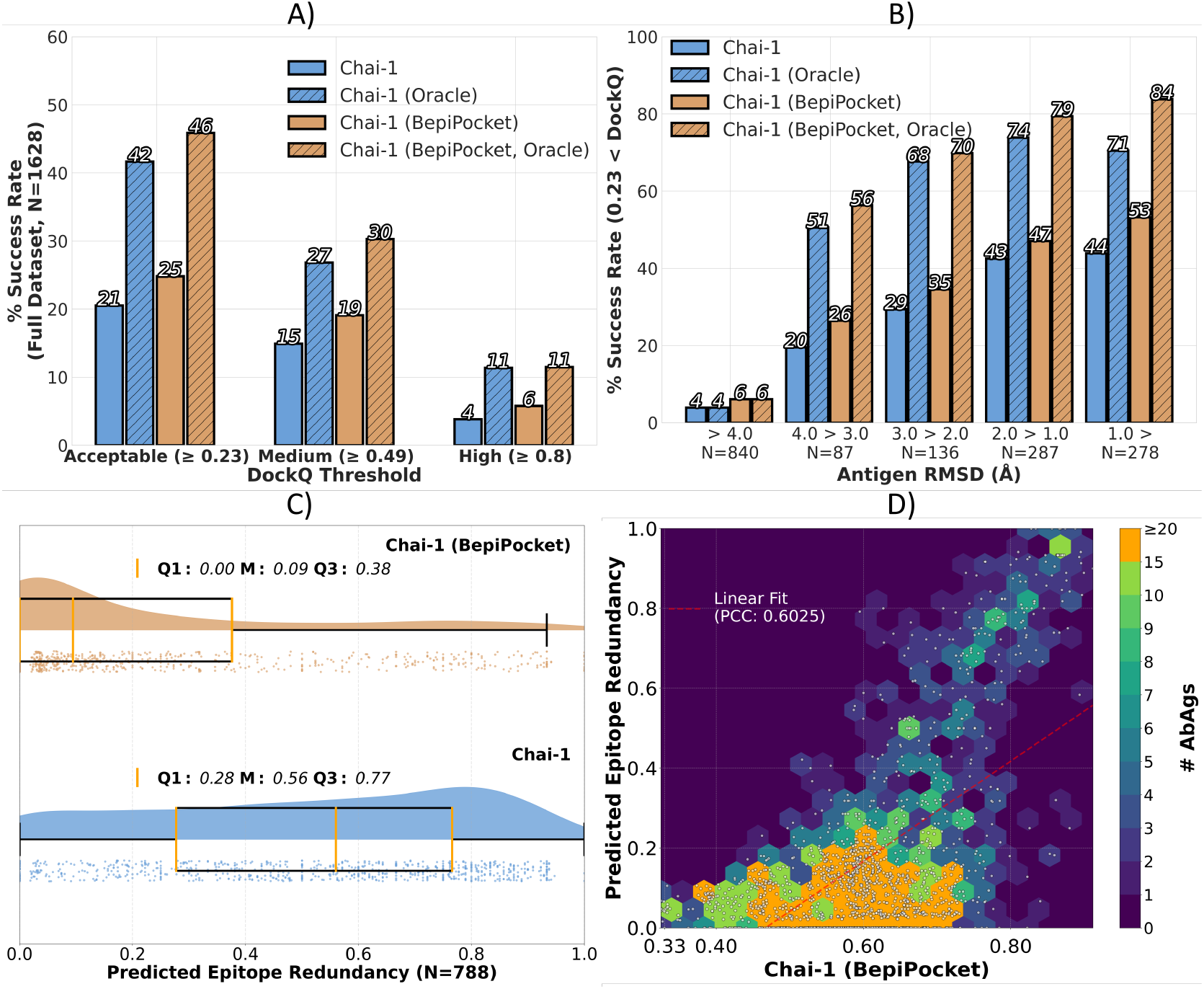
Comparison of Chai-1 AbAg structure prediction using 30 random seeds versus 30 runs with our antibody-epitope pocket restraint algorithm, BepiPocket. **A)** Percentages of AbAg complexes (y-axis, % success rate) with acceptable, medium, or high DockQ accuracy (CAPRI thresholds) using 30 Chai-1 runs with different seeds and 30 BepiPocket runs. Each method creates 150 structures per complex. Results are shown for the highest-confidence structure (Chai-1 or Chai-1 (BepiPocket)) and the best-by-DockQ structure (Oracle). **B)** AbAgs are grouped by antigen structure quality, measured as the best RMSD (Å) among five Chai-1 predicted antigen structures (x-axis), compared to the solved structure. For each group, the percentage of AbAgs with acceptable DockQ accuracy for structures from Chai-1, Chai-1 (BepiPocket), or Oracle selection is shown. **C)** Epitope redundancy of 150 structures predicted using Chai-1 or Chai-1 (BepiPocket) (x-axis) for the 788 AbAgs with antigen RMSD < 4 Å. Each dot is a single AbAg. Higher values indicate greater epitope redundancy. Density plots illustrate the distribution of values, and yellow lines denote Q1, median (M), and Q3. **D)** Median confidence scores of 150 Chai-1 (BepiPocket) predicted structures per AbAg (x-axis) plotted against corresponding epitope redundancy values (y-axis) for 1,628 AbAgs. Each dot represents one AbAg. To illustrate density, dots are overlaid on 2D hexagonal bins colored by the number of AbAgs per bin (color scale capped at 20).

We expected that performance depends on antigen modeling quality. To examine this, AbAgs were grouped into five bins based on the best RMSD (Å) among five Chai-1-predicted antigen structures per AbAg. Within each bin, we examined the percentage of AbAgs obtaining above acceptable CAPRI DockQ quality. For AbAgs with poor antigen modeling quality (RMSD > 4 Å, comprising over half the dataset, n = 840), success rates were low. However, success rates increased with antigen accuracy (**Figure 2B**).

Importantly, the benefit of BepiPocket over Chai-1 with seed variation also increased with antigen quality. While BepiPocket outperformed seed-based Chai-1 runs across all bins, the improvement was particularly pronounced in the most accurate group (RMSD < 1.0 Å), where it achieved a 10% higher success rate, compared to just 2% in the poorest-quality group (RMSD > 4.0 Å; **Figure 2B**).

We reasoned that one factor contributing to BepiPocket’s improved performance is that the method encourages Chai-1 to explore more diverse antibody binding sites. That is, by using variable restraint residues over 30 runs, BepiPocket increases the diversity of docking sites between the antibody and antigen, raising the chance of identifying the correct binding site compared to simply varying random seeds.

**Figure 3** provides one case illustrating this. Here, we show the structure prediction for an antibody targeting Hyaluronidase, a bee venom allergen (PDB 2J88) (27). The ground truth antibody and antigen structures are shown in grey and black, respectively; predicted antibodies are colored blue to red based on Chai-1 confidence. With 30 different seeds, Chai-1 repeatedly places the antibody at the same incorrect site, failing to explore alternatives. In contrast, with BepiPocket many alternative sites are explored, and although some predictions are incorrect, the correct binding site is identified with high confidence.

**Figure 3.**
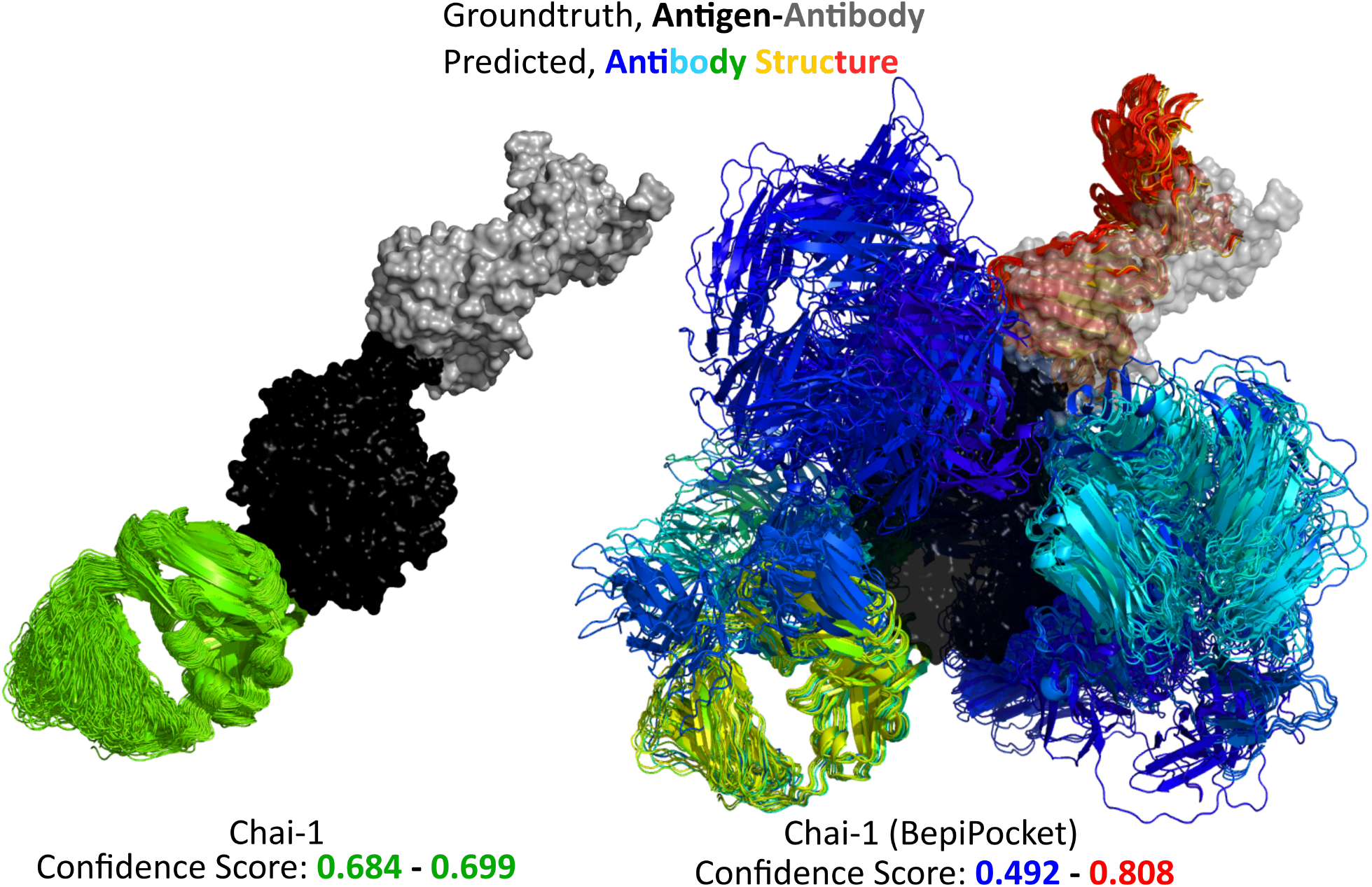
Structure prediction for an antibody targeting Hyaluronidase, a bee venom allergen. The crystal structure of a monoclonal IgG Fab fragment (grey) bound to Hyaluronidase (black) is shown (PDB 2J88). 150 AbAg structures were predicted using Chai-1 with 30 different seeds (left) and Chai-1 (BepiPocket) (right). Predicted antibody structures were superimposed by aligning antigens to the solved antigen structure; predicted antigens were omitted for clarity. For Chai-1 (BepiPocket), five low-confidence predictions were excluded for visualization. Antibodies are colored by Chai-1 confidence (0.8ipTM + 0.2pTM), from low (blue) to high (red). Score ranges are also indicated.

To assess whether BepiPocket generally leads to less redundant epitope predictions, we next measured redundancy of epitope residue sets across different predicted structures for a given AbAg. AgIoU (Antigen Intersection over Union) was used to measure the match between the predicted epitope residue sets of AbAg structure pairs. This metric ranges from 0 (no overlapping epitope residues) to 1 (perfect overlap). For details on this metric and how epitopes residues were defined, refer to Methods. For each AbAg, we computed AgIoU between all unique pairs among the 150 predicted structures, resulting in 11,175 pairwise comparisons per complex. From these values, we then calculated the median AgIoU, reflecting the overall epitope redundancy for that AbAg. We term this metric epitope redundancy, which will be high if all structures are predicted to have the same epitope residue set (low diversity), and low in cases where the structures explore a diverse epitope space.

**Figure 2C** shows results for the 788 AbAgs with antigen RMSD < 4 Å. Visually, BepiPocket predictions are markedly less redundant, with an epitope redundancy value of 0.10 compared to 0.56 for Chai-1 seed-based predictions—a more than fivefold difference. Interestingly, the plot also reveals cases where BepiPocket produces highly redundant epitope predictions (upper right corner of **Figure 2C**). We speculate that this behaviour could be linked to the confidence of the modeled AbAg structures. Each of the 150 predicted structures per AbAg has a confidence score. By plotting the median of these scores against the corresponding per AbAg epitope redundancy, we observed a strong correlation (Pearson’s correlation coefficient, PCC = 0.6025; **Figure 2D**). In practice, this means that when Chai-1 is highly confident, it tends to generate very similar structures across runs, often overriding the imposed pocket restraints and leading to redundant epitope sets. This occurs because the BepiPred-3.0–derived restraints are not enforced as hard constraints and can be ignored by the model. A more detailed analysis is provided in Supplementary S2, which includes the Chai-1 version of Figure 2D and a structural illustration of restraints being bypassed. Importantly, this supplementary analysis also shows that BepiPocket is particularly effective at creating diverse predicted antibody binding sites when Chai-1’s median confidence lies in the 0.6-0.8 range.

In conclusion, antibody-epitope pocket restraints guided by BepiPred-3.0 substantially enhance Chai-1 structure prediction. The BepiPocket algorithm, drives Chai-1 to evaluate a diverse set of antibody binding sites, resulting in less predicted epitope redundancy. Moreover, the benefit of using BepiPocket is linked to antigen modeling quality, indicating that much can be gained by improving the modelled antigen accuracy.

### Multiple sequence alignment improves Chai-1 antibody-antigen structure prediction

In the previous section, we showed that using an antibody-agnostic B-cell epitope predictor—such as BepiPred-3.0—to guide consecutive Chai-1 structure prediction runs improves AbAg prediction accuracy more effectively than simply varying random seeds. This improvement was strongly linked to the structural accuracy of the modelled antigen. Given that a large portion of the dataset contained poorly modelled antigens, we next focused on improving antigen quality by incorporating the use of MSA inputs.

In this analysis, we further included variants of the AlphaFold tool (AlphaFold-2.3 (16) and AlphaFold-3.0 (18)), to evaluate the performance of our Chai-1 based approaches. We did not use structural templates for any method.

For all analyses in this result section, five AbAg structures were made for each method (AlphaFold or Chai-1) and MSA input variation. To compute antigen RMSD, we, as before, predicted 5 unbound (no antibody) antigen structures with each method using a single seed. For antibody RMSD, we used the bound antibodies from the five predicted AbAg structures.

Overall, we found that MSA input had a strong impact on AbAg structural accuracy. AlphaFold-3 (De Novo) performed poorly without MSA —substantially worse than the earlier version AlphaFold-2.3 using ColabFold-generated MSAs (28). In contrast, Chai-1 performed relatively well even without MSA, likely due to its use of ESM2 language model embeddings. Adding MSA input to AlphaFold-3 via its built-in HMMER-based pipeline (29) substantially increased DockQ accuracy. Chai-1 also benefited from this and even a basic MMSeqs2-generated MSAs (30) with the default maximum depth 300 of against the ColabFold’s UniRef30 database, nearly doubled its success rate (Chai-1 (ShortMSA)), making it comparable to AlphaFold-3 (MSA). Increasing MSA depth and sensitivity, and pairing multimeric antigen MSAs by taxonomy, Chai-1 (DeepTaxPair) surpasses AlphaFold-3 (MSA) across all DockQ thresholds. Although the original Chai-1 implementation used jackhmmer, our results suggest that a MMSeqs2 MSA pipeline is sufficient for accurate AbAg structure prediction (**Figure 4A**). We also examined antigen accuracy measuring the atomic distance in Å RMSD between predicted and solved antigens, across all AbAgs. Accuracy was substantially improved when comparing Chai-1 predictions without MSA to those with MSA input (**Figure 4B**), while antibody structure quality remained largely unaffected (Supplementary Figure S3A). However, comparing Chai-1 (DeepTaxMSA) to Chai-1 (ShortMSA), antigen quality only improved slightly in the median. Also, when evaluating the impact of taxonomy-based pairing on the 321 AbAgs with multimeric antigens—where this strategy could apply—we observed no improvement in antigen accuracy and only minimal gain in AbAg accuracy (see Supplementary Section S3).

**Figure 4.**
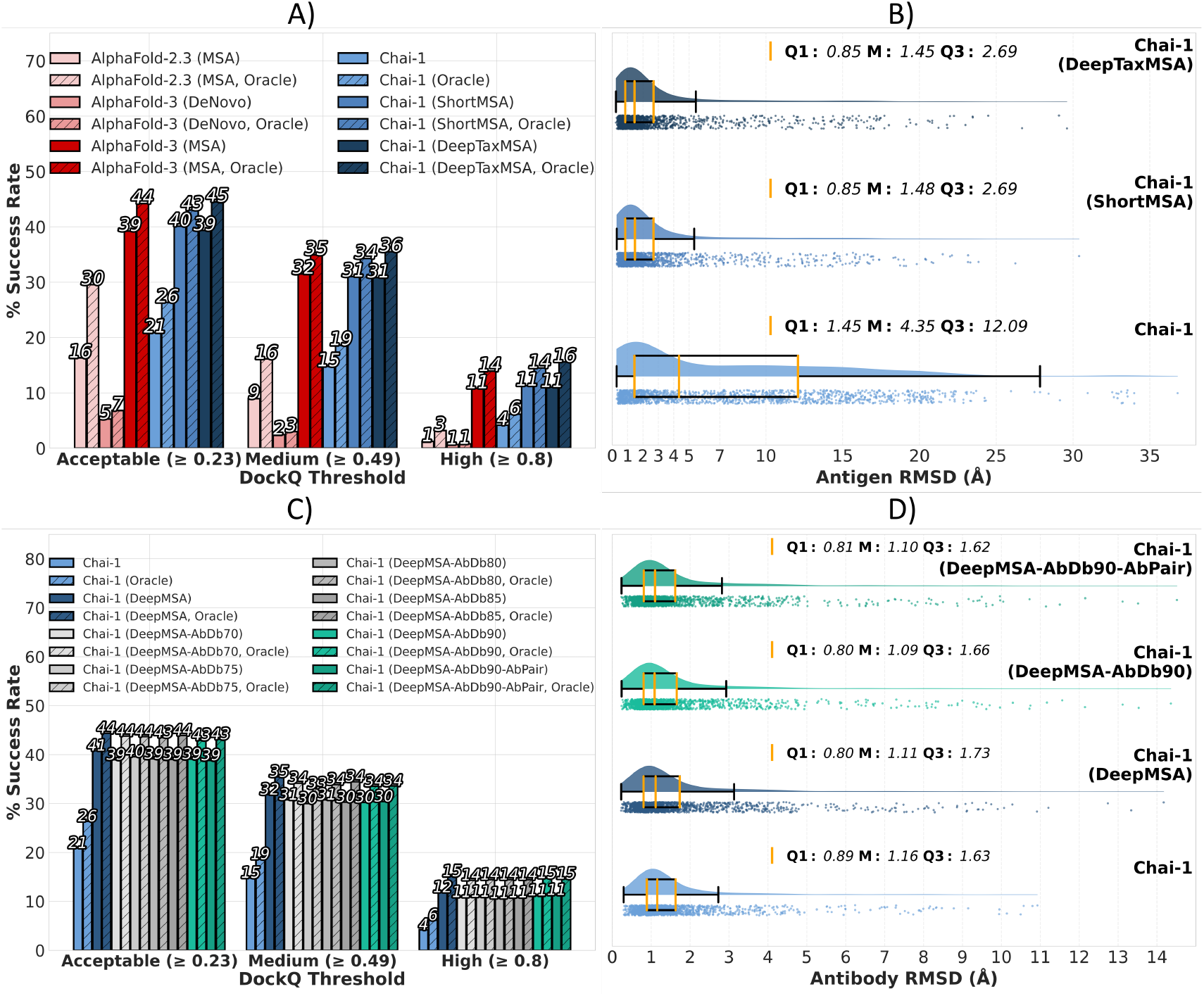
Comparison of AlphaFold and Chai-1 AbAg structure prediction across 1,628 complexes using different MSA inputs. **A)** Bar plot showing the percentage of AbAg predictions meeting CAPRI-standard DockQ thresholds—acceptable (≥ 0.23), medium (≥ 0.49), and high (≥ 0.80)—referred to as success rate (y-axis). Each method produced 5 structures per AbAg; shown are the highest-confidence prediction (AlphaFold or Chai-1) and the best-by-DockQ prediction (Oracle). Methods include: AlphaFold-2.3 (MSA), AlphaFold-3 (De Novo), AlphaFold-3 (MSA), Chai-1, Chai-1 (ShortMSA), and Chai-1 (DeepTaxMSA). **B)** Distribution of antigen RMSD (Å, x-axis) for the Chai-1 variants in A (y-axis). For each AbAg, the best (lowest RMSD) of 5 predicted antigen structures is shown using rain-cloud and box plots. Yellow lines indicate Q1, median (M), and Q3. **C)** As in A), comparing Chai-1 without MSA or with UNIREF30 MSA input to variants using paired antibody sequence databases for antibody MSA generation. ‘X’ in ‘AbDbX’ indicates the MMSeqs2 identity threshold used for redundancy reduction in these databases. The variant testing light/heavy chain MSA pairing is highlighted in teal. **D)** As in B), but instead comparing antibody RMSD values (Å, x-axis) for each AbAg for methods: Chai-1, Chai-1 (DeepMSA), Chai-1 (DeepMSA-AbDb90), and Chai-1 (DeepMSA-AbDb90-AbPair).

Since the observed gains were driven by improved antigen modelling, we next tested whether optimizing antibody modelling could also enhance AbAg structure accuracy. Specifically, we assessed whether refining the antibody MSA input would improve the quality of the antibody structure and thus the full complex. Instead of using UniRef30, we built antibody-specific sequence databases from paired heavy and light chains in the SABDAB and OAS datasets (31, 32), and generated MSAs using MMSeqs2 with various identity thresholds from 70% to 90% (for details on this, see Methods). We did not use taxonomy-based pairing of antigen MSAs and so for a direct comparison re-did the structure prediction of Chai-1 (DeepTaxPair), but without the taxonomy pairing. We called this method (Chai-1 DeepMSA). We found that the custom antibody MSAs did not improve AbAg structure prediction. We also tested whether pairing light and heavy chain sequences in the MSAs (Chai-1 DeepMSA-AbDb90-AbPair) improved performance relative to unpaired MSAs (Chai-1 DeepMSA-AbDb90), but found no gain (**Figure 4C**). In terms of antibody RMSD across all predicted AbAgs, we observed only very slight improvements in median (**Figure 4D**).

### BepiPred-3.0 and DiscoTope-3.0 guided pocket restraints combined with multiple sequence alignment input further improve accuracy

In the previous section, we showed that adding MSA input to Chai-1 substantially improved antigen modelling accuracy, which in turn enhanced AbAg complex predictions. Earlier, we also demonstrated that the benefit of BepiPred-3.0–guided pocket restraints (without MSA input) was strongly tied to antigen quality. Given this, we revisited our pocket selection strategy from the second results section, but now using antigens modelled with the DeepTaxMSA method. For brevity, call this method ‘MSA’.

With the added MSA input, we applied the same approach as before, predicting structures for 1,628 AbAgs in our dataset with Chai-1, using 30 different seeds (Chai-1 MSA) or 30 iterations using the BepiPocket algorithm (Chai-1 MSA-BepiPocket). To compute the antigen RMSD, we again predicted 5 unbound (no antibody) antigen structures with Chai-1 using a single seed.

The overall conclusions mirror those from the second results section, but with higher accuracy due to the added MSA input. BepiPocket improved performance, with 48%, 40% and 12% of AbAgs reaching acceptable, medium, and high DockQ (CAPRI thresholds), compared to 42%, 33%, and 12% for Chai-1 (MSA) (**Figure 5A**). When restricting to the 788 AbAgs with antigen RMSD < 4 Å, BepiPocket predictions were markedly less redundant than those from Chai-1 with random seeds, achieving an epitope redundancy value of 0.06 versus 0.62, a more than four fold difference (**Figure 5B**).

**Figure 5.**
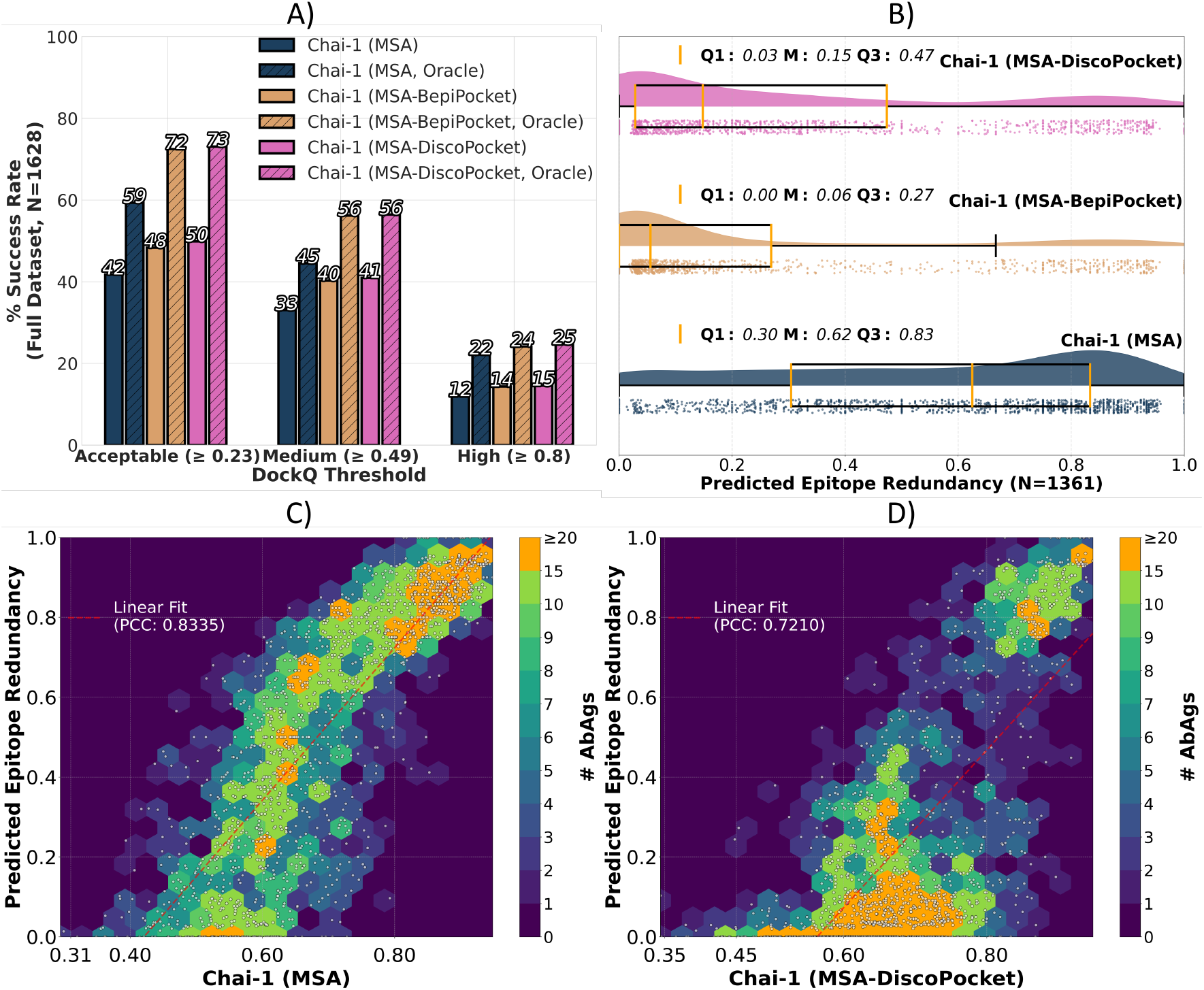
Comparison of Chai-1 AbAg structure prediction using 30 random seeds versus 30 runs to our antibody-epitope pocket restraint algorithms. **A)** Percentages of AbAg complexes (y-axis, % success rate) with acceptable, medium, or high DockQ accuracy (CAPRI thresholds) using 30 Chai-1 runs with different seeds and 30 BepiPocket or DiscoPocket runs. Each method creates 150 structures per complex. Results are shown for the highest-confidence structure (using Chai-1’s confidence score) and the best-by-DockQ structure (Oracle). **B)** Epitope redundancy of 150 structures predicted using Chai-1, BepiPocket or DiscoPocket (x-axis) for the 788 AbAgs with antigen RMSD < 4 Å. Each dot is a single AbAg. Higher values indicate greater epitope redundancy. Density plots illustrate the distribution of values, and yellow lines denote Q1, median (M), and Q3. **C)** Median confidence scores of 150 Chai-1 (MSA) predicted structures per AbAg (x-axis) plotted against corresponding epitope redundancy values (y-axis) for 1,628 AbAgs. Each dot is one AbAg. To illustrate density, dots are overlaid on 2D hexagonal bins colored by the number of AbAgs per bin (color scale capped at 20). **D)** Same as C), but with Chai-1 (MSA-DiscoPocket) predicted structures.

We observed that with the large increase in modelled antigen accuracy, DiscoTope-3.0 was now more capable than BepiPred-3.0 at incorporating epitope contacts found in solved structures as pocket restraints (for details, see Supplementary S5).

To investigate how this impacted the actual performance gains, we re-did the pocket restraint guided structure prediction using DiscoPocket instead. DiscoPocket further improved performance across all DockQ CAPRI thresholds, with 50%, 41%, and 15% of AbAgs having acceptable, medium, and high quality (**Figure 5A**). Similar to BepiPocket, DisoPocket also predicted markedly less redundant epitopes (**Figure 5B**) Plotting the median of the 150 confidence scores per AbAg against redundancy showed a strong correlation (PCC = 0.8335) for Chai-1 (MSA), but a substantially lower correlation (PCC = 0.7210) for DiscoPocket (**Figure 5C, D**). This indicates that the tendency of highconfidence predictions to collapse into redundant epitopes is mitigated when pocket restraints are guided by DiscoTope-3.0. This effect is especially clear in the density of AbAgs with confidence scores of 0.6-0.8, where DiscoPocket yields consistently low redundancy, thus expanding the high structural diversity to also cover the intermediate confidence range. For both BepiPocket and DiscoPocket, the performance gain over Chai-1 remained closely linked to antigen RMSD, particularly pronounced for highly accurate antigens (RMSD < 1.0 Å), where 13 and 15% higher success rate were achieved. For this plot and BepiPocket version of Figure 5D, see Supplementary section S6.

In conclusion, the combination of MSA input and BepiPred-3.0 or DiscoTope-3.0–guided pocket restraints substantially enhances Chai-1 structure prediction.

### BepiPocket and DiscoPocket performance gains are robust on independent test data

The training data for both Chai-1, BepiPred-3.0 and DiscoTope-3.0 consists of previously solved AbAg structures from the PDB. Such data might have been included in the training of Bepipred, Discotope, AlphaFold and Chai-1 methods thus potentially imposing a bias in our performance evaluations. To investigate the impact of this, we evaluated our methods on AbAgs released after the training cutoffs of BepiPred-3.0 and DiscoTope-3.0 (29 September 2021) as well as Chai-1 (12 January 2021).

That is, from the full dataset of 1,628 AbAgs, we constructed two subsets. The “After” subset reuses the independent test set from Clifford et al. (17): 109 AbAgs released after 30 September 2021, filtered to ensure ≤65% antigen and ≤95% antibody MMseqs2 identity to any pre-cutoff AbAg. After removing complexes exceeding the 2,048-residue limit of Chai-1, 105 AbAgs remained. The “Before” subset consisted of the remaining 1,529 AbAgs. We find that AbAg prediction accuracy is substantially lower on the independent ‘After’ set than on the ‘Before’ set. Importantly, however, the relative performance gain from using epitope prediction tools to guide AbAg structure prediction remains consistent across both subsets (**Figure 6A**).

**Figure 6.**
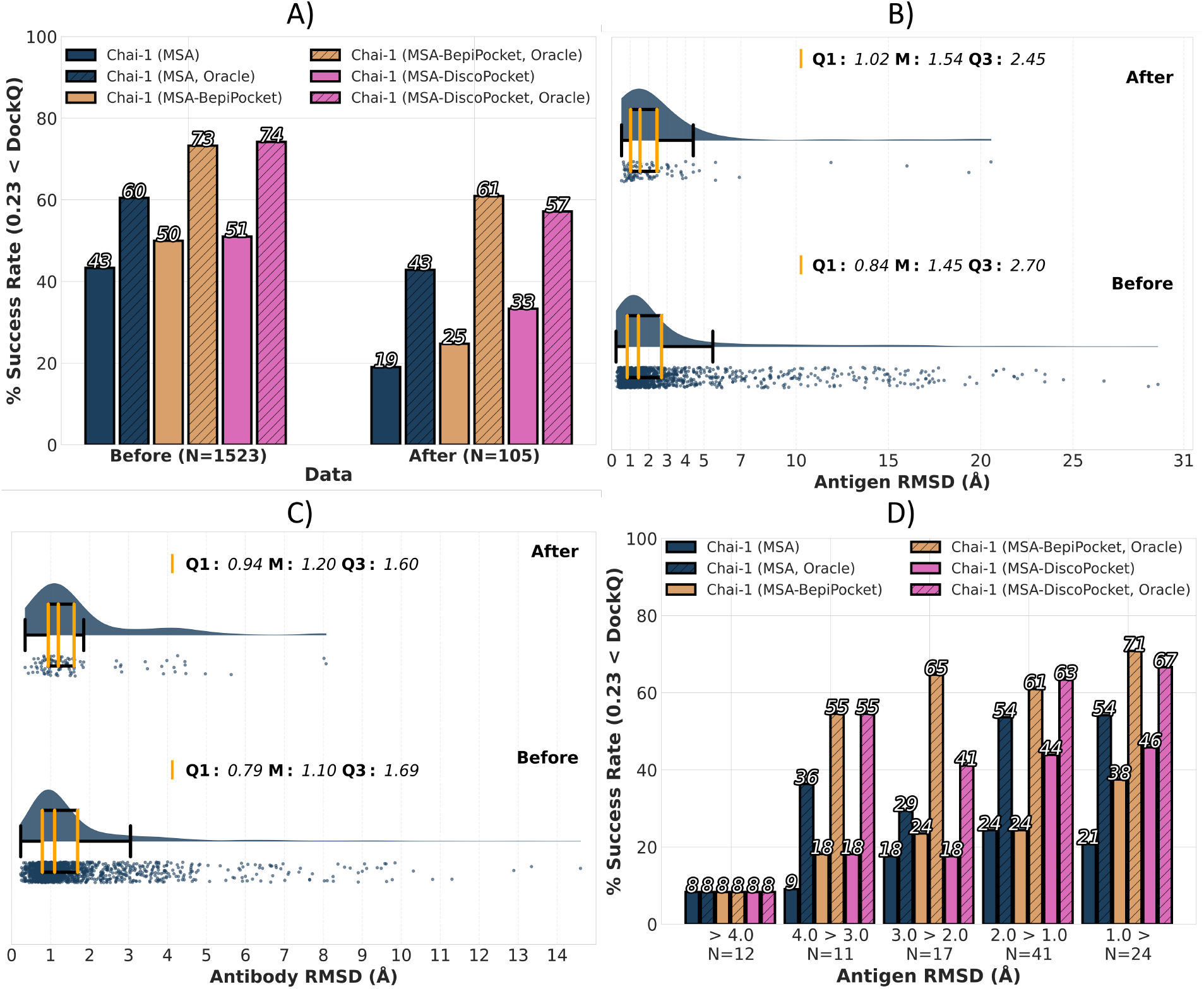
Performance of Chai-1 with and without BepiPocket or DiscoPocket on AbAg subsets released before and after training cutoffs. **A)** Percentage of AbAg complexes (y-axis, success rate) in the Before and After subsets achieving acceptable DockQ accuracy (>0.23). Each method was run with 30 iterations (30 random seeds, DiscoPocket runs, or BepiPocket runs), creating 150 structures per complex. Results are reported for the highest-confidence prediction and for the best structure per complex ranked by true DockQ (Oracle). **B)** Distribution of unbound antigen RMSD (Å, x-axis) for Chai-1 MSA in the Before and After subsets. For each AbAg, the best (lowest RMSD) of 5 predicted antigen structures is shown using raincloud and box plots. Yellow lines indicate Q1, median (M), and Q3. **C)** As in B), but comparing antibody RMSD values (Å, x-axis). **D)** AbAgs in the After subset were grouped by the antigen structure quality measurements displayed in B). For each group, we show the percentage of AbAgs with acceptable DockQ accuracy for structures from Chai-1 (MSA), Chai-1 (MSA-DiscoPocket), or Oracle selection.

To investigate the source of this performance decrease, we repeated the antigen and antibody RMSD analysis from our third results section, this time focusing only on Chai-1 (MSA) and evaluating the two subsets separately rather than the full dataset. As expected, RMSD is worse on AbAgs released after the cutoff, with both antigen and antibody RMSD increasing by a median of around 0.1 Å (**Figure 6B, C**). This indicates that part of the performance drop can be attributed to worse antigen and antibody modeling. However, the increase in RMSD is small compared to the overall drop in complex accuracy. For example, 54% of AbAgs (57/105) in the ‘After’ subset still achieve Ag and Ab RMSD values below 3 Å and 1.5Å, respectively, which means that in many cases the antigen and antibody are modeled well individually, but Chai-1 still fails to position the antibody correctly on the antigen.

Nonetheless, because our pocket-based approaches are strongly dependent on modeled antigen quality—and this does not differ dramatically between the ‘Before’ and ‘After’ subsets— we observe the same trends demonstrated earlier on the After compared to the Before data sets. Specifically, BepiPocket and DiscoPocket improve accuracy when the antigen can be modeled well and its performance gains are robust also in more challenging post-cutoff benchmarks (**Figure 6D**).

## Discussion

Here, we have demonstrated that integrating B-cell epitope prediction tools to guide pocket restraints for Chai-1 substantially improves antibody–antigen (AbAg) complex prediction. By using sequence-based or structure-based epitope predictors BepiPred-3.0 and DiscoTope-3.0 to select epitope residues for antibody-epitope restraints, more accurate and diverse antibody binding site predictions are produced compared to Chai-1 with random seed variation. We call these approaches BepiPocket and DiscoPocket, respectively.

The effectiveness of this strategy strongly depends on antigen structural quality: performance gains were substantially higher when the antigen could be modeled accurately. If the predicted antigen structure is incorrect or inconsistent, correct placement of the antibody becomes impossible because the true epitope is not there. To improve antigen modeling, we generated multiple sequence alignments (MSAs), which substantially enhanced predicted antigen and AbAg accuracy—even simple MMseqs2-generated MSAs (ShortMSA) roughly doubled performance. For dimeric antigens, pairing MSAs by taxonomy showed little benefit, likely because species-level cooccurrence does not guarantee interaction and paired sequences were sparse relative to MSA depth. This aligns with observations from Hopf et al. (2014) (33), showing that naïve taxonomic pairing of sequences can introduce false positives, since species co-occurrence does not guarantee conserved interaction, and insufficient alignment depth further limits reliable co-evolutionary signal. More advanced strategies, such as using orthologs, interaction databases, or genomic context, may better capture co-evolutionary signals.

Custom antibody MSAs derived from paired heavy and light chains databases, redundancy reduced at various sequence identity thresholds, had very little impact on AbAg accuracy or antibody RMSD, suggesting that antibody MSAs provide limited benefit for Chai-1 predictions. These findings corroborate other studies showing that antibody structure prediction tools not relying on MSAs can perform as well as, or even better than, MSA-based methods (34–36).

Benchmarking on AbAg structures released after the Chai-1, BepiPred-3.0 and DiscoTope-3.0 training cutoffs revealed a pronounced performance drop, with accuracy nearly halved on the independent “After” test set compared to newer structures. Importantly, however, the relative performance gain from using BepiPocket or DiscoPocket remained consistent across both subsets, highlighting that epitope-guided restraints provide a robust advantage even in harder-to-predict cases.

An issue with both BepiPocket and DiscoPocket is that very high-confidence Chai-1 predictions can override antibody-epitope pocket restraints. In this current work, only single residue–antibody restraints have been applied. One potential solution would be to switch to predicted epitope patches or regions, applying multiple antigen residues simultaneously to better enforce binding site guidance. Additional strategies, such as specifying pairwise contacts between HCDR3 residues and selected epitope residues, may further improve restraint effectiveness. Finally, for computational efficiency, predictions could track which restraints have already been satisfied and prioritize predictions covering previously unrestrained epitope sites.

In summary, BepiPocket and DiscoPocket are simple yet highly effective approaches for improving predicted antibody–antigen accuracy. They provide a strong baseline and point to a promising direction for more accurate in silico prediction of antibody targets, with major applications in diagnostics and therapeutic antibody development. We anticipate that more advanced algorithms, incorporating epitope patches and pairwise contacts, could further enhance performance. Both tools are available as a standalone package, making them accessible to experts and non-experts alike.

## Methods

### Structural data

We first extracted all solved AbAgs from our previous study, Clifford et al. (17). This included a redundancy reduced dataset of 1733 antibody-antigen structures with a resolution lower than 3.5 Å and an R-factor below 0.26, deposited to the Protein Data Bank (PDB) biological assembly FTP server before February 2, 2023. Each solved structure consisted of one antibody containing both light and heavy chain variable domains, as well as one non-antibody (antigen) protein chain with at least 40 residues. Next, we removed all structures whose total number of residues exceeded the 2048 residue limit for Chai-1 structure prediction, leaving 1628 remaining structures. Among these, 1,307 had monomeric antigens and 321 had dimeric antigens.

### Antibody-Antigen structure prediction

Structure predictions for all 1,628 experimentally solved antibody–antigen (AbAg) complexes were performed using AlphaFold-2.3, AlphaFold-3, and Chai-1. Each of these tools typically requires three types of input: protein sequences, multiple sequence alignments (MSAs) of related sequences, and optionally, structural templates. We did not provide template structures for any predictions.

### AlphaFold Antibody-Antigen structure prediction

For AlphaFold-2.3, we used the ColabFold MMSeqs2 pipeline to generate MSAs for all antibody and antigen sequences, using UniRef30 as the target database. This version of UniRef30 is a 30% sequence identity–clustered version of UniRef100, publicly distributed by the Steinegger lab at https://colabfold.mmseqs.com/. This is also the default database used by ColabFold. These predictions are referred to as AlphaFold-2.3 (MSA). For AlphaFold-3, we first predicted structures without any MSA input, which we refer to as AlphaFold-3 (De Novo). We then also used AlphaFold-3’s recommended inference pipeline, which uses the HMMER version 3.4 jackhmmer, to generate unpaired MSAs and automatically performs MSA pairing; this is referred to as AlphaFold-3 (MSA).

### Chai-1 Antibody-Antigen structure prediction

Chai-1 predictions were performed using version 0.6.0 installed via pip. Initially, we ran structure predictions without providing any MSA input, relying solely on the ESM2 protein language model embeddings that Chai-1 generates internally from input sequences. This mode is referred to as Chai-1.

We used MMSeqs2 to generate antibody and antigen MSA inputs for Chai-1 structure predictions. In one approach, MSAs were constructed using default MMSeqs2 settings, with a sensitivity of 5.6 and a maximum alignment depth of 300 sequences, against the UniRef30 database. These MSAs were used as input to Chai-1, and the resulting prediction mode is referred to as Chai-1 (ShortMSA).

In a second approach, we increased the depth and sensitivity of MSAs by setting MMSeqs2 sensitivity to its maximum value of 8 and raising the maximum alignment depth to 10,000 sequences. In addition, we leveraged Chai-1’s support for user-defined pairing keys. As described in Chai-1’s documentation, a pairing key is a string that specifies how MSAs from different chains should be grouped and aligned during structure prediction. Pairing is performed across all sequences sharing the same key. Although this key is typically a species identifier, any shared attribute can be used to define the grouping. To enable taxonomy-based pairing for our AbAg dataset, we mapped each antigen sequence to its corresponding NCBI taxonomy ID using UniRef30 taxonomy annotations. These taxonomy IDs were then assigned as pairing keys, allowing Chai-1 to pair antigen sequences originating from the same species. Structure predictions generated with these taxonomically paired MSA inputs are referred to as Chai-1 (DeepTaxMSA). We also used a third mode, with the same settings, but without the taxonomical pairing, which we called Chai-1 (DeepMSA).

Finally, we evaluated whether antibody-specific MSAs could improve prediction accuracy by creating paired antibody sequence databases. To do this, we collected all paired light–heavy chain variable domain sequences from SABDAB (as of February 5, 2025), resulting in 13,077 unique pairs. We then extracted an additional 3,086,358 paired sequences from the Observed Antibody Space (OAS). After removing exact duplicates—defined as pairs with identical light and heavy chains— we retained 2,595,991 unique paired antibody sequences. To reduce redundancy within these databases, we concatenated each light–heavy pair and clustered them using MMSeqs2 linear clustering. We applied coverage mode 5 with a minimum coverage of 0.1, and tested a range of MMSeqs2 sequence identity thresholds: 70%, 75%, 80%, 85%, and 90%. After clustering, representative sequences were split back into separate light and heavy chains to form paired MSA databases. For each of these identity thresholds, we generated antibody MSAs using MMSeqs2 with the same high-sensitivity settings as in the DeepMSA pipeline (sensitivity 8, depth 10,000). These antibody MSAs combined with antigen MSAs from the DeepMSA pipeline (against UniRef30) were used for Chai-1 structure predictions, and the corresponding prediction modes are referred to as Chai-1 (DeepMSA-AbDbX), where ‘X’ denotes the sequence identity threshold used for clustering.

### Chai-1 BepiPocket and DiscoPocket Antibody-Antigen structure prediction

We developed a Chai-1 pocket restraint mode for antibody–antigen complexes (AbAgs), guided by epitope likelihood scores generated using the antibody-agnostic B-cell epitope predictors BepiPred-3.0 and DiscoTope-3.0. BepiPred-3.0 is a sequence-based tool that uses a feedforward neural network trained on ESM-2 protein language model embeddings of high-quality solved structures to generate per-residue B-cell epitope scores. ESM2 is a language model developed by Meta’s Fundamental AI Research (FAIR), using a masked language model training approach on 60M protein sequences from the UniRef50 database. DiscoTope-3.0 is a structure-based tool that uses similar data, but instead applies an XGBoost classifier trained on ESM-IF1 inverse folding embeddings (37). Inverse folding seeks to recover the amino acid sequence compatible with a given protein’s three-dimensional structure. ESMIF1 is also a deep learning model developed by FAIR, trained on a large dataset of 12 million AlphaFold and 16,000 CATH structures (38).

This Chai-1 structure prediction mode accepts a ranked list of antigen residues, sorted by predicted epitope likelihood (high to low), as input. For the first run, no restraint is used, allowing Chai-1 to run without any restraint. But then, for each Chai-1 run, the next residue in the ranked list is used as a spatial restraint, enforcing a 10 Å distance to the antibody. All runs were conducted with a fixed seed value of 0.

For BepiPred-3.0, we generated ranked residue lists by applying the predictor to the antigen sequences of all 1,628 AbAgs in our dataset. For DiscoTope-3.0, which requires structure-based input, we first predicted 5 unbound antigen structures using Chai-1 (DeepTaxMSA), and then used the structure with the highest Chai-1 confidence score as input to DiscoTope-3.0.

We refer to these two approaches as BepiPocket and DiscoPocket, respectively. BepiPocket and DiscoPocket were both run for a total 30 runs (1 initial run with no restraint + 29 epitope likelihood restraint based runs).

### Evaluation and performance metrics

We used DockQ version 1 with default options to compute DockQ scores for all predicted antigen– antibody (AbAg) structures generated by either Chai-1 or AlphaFold. DockQ scores evaluate the quality of predicted AbAg interfaces by combining interface RMSD (iRMSD), ligand RMSD (LRMSD), and the fraction of native contacts (Fnat) into a single score ranging from 0 to 1.

To evaluate the accuracy of individual antigen structures, we computed per-chain RMSD values by re-running structure prediction for each Chai-1 mode using only the unbound antigens of the 1,628 AbAgs. For each unbound run, a single run was with seed value 0, generating five antigen per AbAg. RMSD values (in Ångströms) were computed by comparing the predicted and corresponding experimentally solved structures, using only alpha-carbon (Cα) atoms. Antibody RMSD was obtained by comparing antibodies from predicted AbAgs, to corresponding bound antibodies found in solved structures. The final antibody RMSD was obtained by averaging the RMSD of the light and heavy chains. Structures were superimposed using Biopython’s Superimposer, and RMSD was calculated on a per-chain basis. For dimeric antigens, we averaged the RMSD across both antigen chains.

To measure the epitope redundancy between any pair of structures predicted for the same AbAg, we used a metric called AgIoU (Antigen Intersection over Union), used in our previous study Clifford et. al. (17). This metric measures the match between epitope residues, which is defined as any residue on the antigen with one or more heavy atoms within 4 Å of the light or heavy chain. The epitope match between two structures is then measured as the intersection divided by the union of epitopes residues (eq. 1). The code for computing these metrics is available at the following GitHub repository https://github.com/mnielLab/AbAgIoU.git.

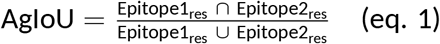

For each AbAg, we computed aggregated epitope redundancy scores, by first computing AgIoU between all unique pairs of predicted structures and then calculating the median of these values. We call this metric ‘epitope redundancy’.

