## Supplementary for "Pocket Restraints Guided by B-Cell Epitope Prediction Improves Chai-1 Antibody-Antigen Structure Modeling"

#### S1: Evaluating Pocket Restraint Selection Strategies for Chai-1 AbAg structure prediction

To evaluate the effectiveness of BepiPred-3.0 for selecting informative pocket restraints, we conducted a preliminary analysis comparing it to random scoring and two structure-based alternatives: DSSP and DiscoTope-3.0. Specifically, we assessed how quickly the top true epitope contact (1Epi, defined as the antigen residue with most antibody contacts; see Figure 1 in Main) is included as a restraint across multiple runs. To enable structure-based scoring, we predicted five unbound (no antibody) antigen structures per complex using Chai-1 with a single seed, and used the one with the highest confidence for scoring.

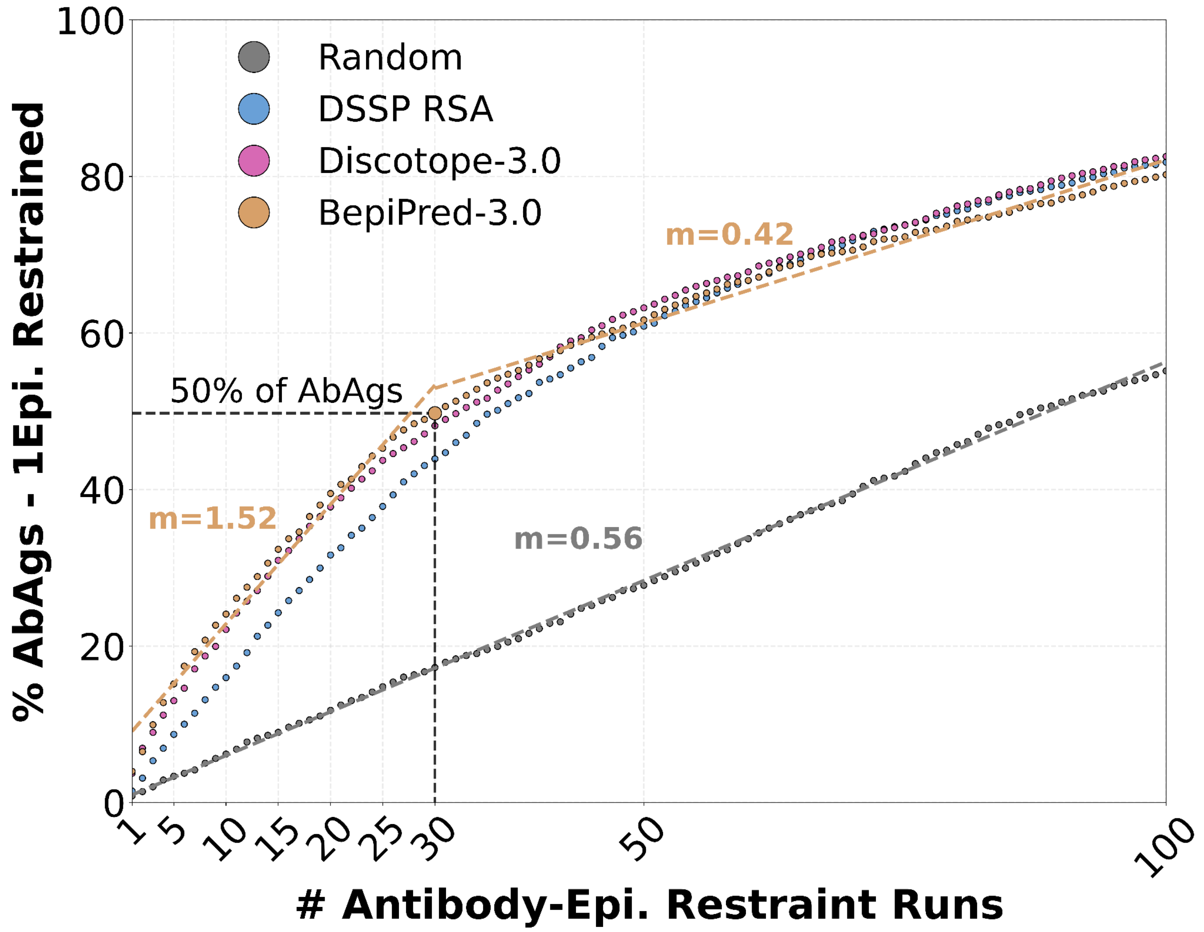


**Fig. S1: Evaluation of epitope restraint selection without AbAg structure prediction.** Percentages of 1,628 AbAg complexes (y-axis) in which the top true epitope contact (i.e., the residue with the most antibody contacts in the solved structure, 1Epi) is included in the restraint, as a function of run number (x-axis). In each run, a new top-ranked residue is selected from a precomputed list, progressively moving down the ranked order. Rankings were derived from BepiPred-3.0 (sequence-based), DiscoTope-3.0, and DSSP solvent accessibility, computed on the highest-confidence Chai-1-predicted antigen structure (from five per complex). A random ranking serves as a baseline. Linear slopes (m) are shown for BepiPred-3.0 (runs 1–30 and 30–100) and Random (runs 1–100).

**Figure S1** shows the percentage of AbAgs (y-axis) in which the 1Epi epitope residue is included as a restraint, as a function of the number of runs (x-axis). Using BepiPred-3.0 to rank residues, approximately 50% (49.75%) of AbAgs include 1Epi by run 30. For comparison, DiscoTope-3.0 achieves 48.16%, and DSSP achieves 43.92%. DSSP RSA scores were obtained by normalizing absolute DSSP values using the maximum residue-specific theoretical values defined by Tien et al. (26). All methods significantly outperformed the random baseline (17.26%). Notably, we observe that the benefit of using BepiPred-3.0 appears to plateau after ~30 runs. That is, the rate of identifying new top epitope contacts follows a linear trend with a high slope (m = 1.52) from runs 1–30, but drops substantially between runs 30–100 (m = 0.42), even falling below the slope for random ranking (m = 0.56).

**S2: BepiPocket decreases redundancy of predicted epitopes, but its restraints are often disregarded by Chai-1 when conflicting with high-confidence predictions**

We investigate epitope redundancy of Chai-1 predicted AbAg structures and the corresponding Chai-1 confidence scoring. For each of the 1628 AbAgs in our dataset, 150 antibody-antigen structures with corresponding confidence scores were predicted using Chai-1 and BepiPocket (Chai-1 with Bepipred guided restraints). For each AbAg, we plotted its epitope redundancy value against the median of these confidence scores **(Figure S2A and B).**

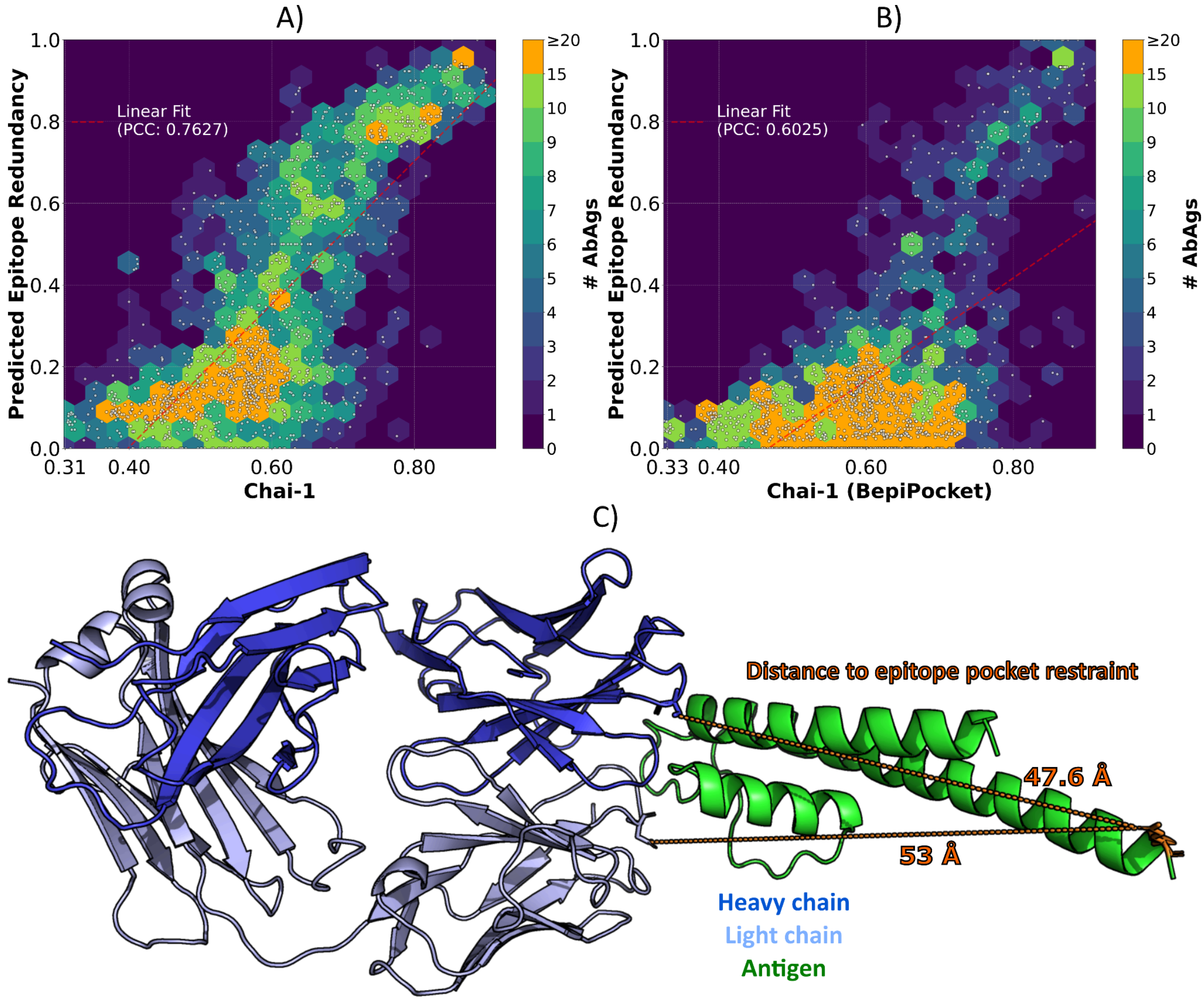


**Fig. S2: The relationship between redundancy of predicted epitopes and Chai-1 confidence scoring. A)** Median confidence scores of 150 Chai-1-predicted structures per AbAg (x-axis) plotted against corresponding epitope redundancy values (y-axis) for 1,628 AbAgs. Each dot represents one AbAg. To illustrate density, dots are overlaid on 2D hexagonal bins colored by the number of AbAgs per bin (color scale capped at 20). **B)** Same as A, but predictions were generated using Chai-1 (BepiPocket). **C)** Example Chai-1 (BepiPocket) prediction for an antibody targeting KcsA (potassium channel from Streptomyces, PDB 7M2J), illustrating a case where the pocket restraint is not maintained. The closest distances from the specified restraint residue to the heavy and light chains are 47.6 Å and 53 Å, respectively (indicated in orange).

For both methods, there is a strong linear correlation between the confidence score and the epitope redundancy score with Pearson's correlation coefficients (PCCs) of 0.7627 and 0.6025 for Chai-1 and Chai-1 (BepiPocket), respectively. This indicates that if Chai-1 is more confident in its structure predictions, predicted epitopes are more likely to be redundant. For BepiPocket this correlation is reduced, indicating that the epitope redundancy is less affected by the model confidence, and most of the upper right corner is shifted to be below 0.2 predicted epitope redundancy compared to Chai-1. Visually, it is also clear that BepiPocket is particularly effective at creating diverse predicted antibody binding sites when the median confidence lies in the 0.6-0.8 range.

When we further investigated these highly confident BepiPocket structure predictions with high epitope redundancy (AbAgs in the upper right corner of Figure S2B), we found that in these cases, the set epitope pocket restraints are ignored. In **Figure S2C,** we showcase an example of the structure of one such antibody-antigen complex, PDB: 7M2J (Chains: A, B and C). The confidence scores for the 150 structures generated for this AbAg range from 0.84-0.87, which is very high, as only (0.61% (10/1628) of our studied AbAgs received a confidence score equal or above 0.84. For this AbAg structure prediction and others receiving the same confidence, we found many instances where epitope pocket restraints were ignored.

### S3: Taxonomy pairing impact on the quality of predicted multimeric antigens

Taxonomy-based pairing of antigen MSAs for AbAgs with multimeric antigens is only relevant when sequences across the antigen chains share the same taxonomy ID. To assess whether this holds for our dataset, we examined the MSAs generated with MMSeqs2 against the UNIREF30 database, using default parameters with a sensitivity of 8.0 and a maximum depth of 10,000. Among the 321 AbAgs with multimeric antigens (all dimeric), we evaluated whether their antigen MSAs had sequences with matching taxonomy IDs. For 29 cases, no such matches occurred, and in the most cases (158), shared taxonomy IDs occurred between 1 and 100 times (**Figure S3A**). This suggests that taxonomy-based MSA pairing is applicable in a subset of multimeric cases and could influence the modelling accuracy of multimeric antigens. To investigate this, we ran Chai-1 structure prediction with a single seed for these MSAs, generating 5 structures for each antigen, with (DeepTaxMSA) and without taxonomy pairing (DeepMSA). We found that taxonomy pairing has little impact on multimer antigen accuracy, comparing Chai-1 (DeepMSA) and Chai-1 (DeepTaxMSA) (**Figure S3B**). We did the same structure prediction on the full AbAgs with multimer antigens, which showed slight improvements in DockQ accuracy across all CAPRI quality thresholds (**Figure S3C**).

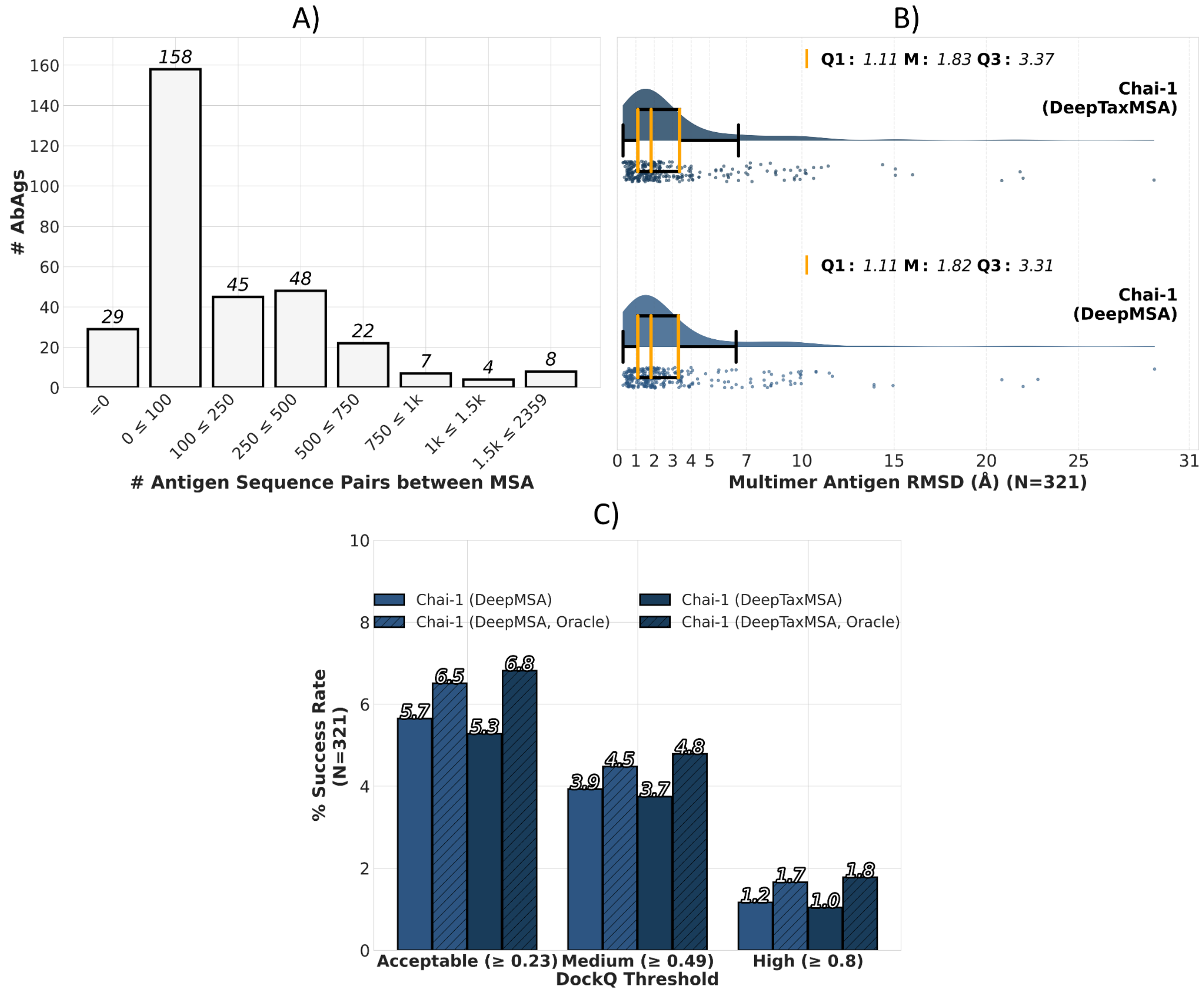


**Fig S3: Modelled accuracy of the 321 antibody-antigen complexes with multimeric antigens, when using taxonomy to pair sequences in MSAs. A)** Counts of AbAgs with multimeric antigens (y-axis), grouped according to how many sequences across antigen MSA had identical taxonomy IDs (x-axis). **B)** Multimer antigen RMSDs for all 321 AbAgs is displayed as a rain plot (Å, x-axis). For each AbAg, the best (lowest RMSD) of 5 antigen structures predicted using Chai-1 (DeepMSA) or Chai-1 (DeepTaxMSA), are shown as dots. Density and boxplots indicate distribution. Yellow lines indicate Q1, median (M), and Q3. **C)** Bar plot showing the percentage of multimer AbAg structure predictions using Chai-1 (MSA) or Chai-1 (DeepTaxMSA), surpassing CAPRI-standard DockQ accuracy thresholds—acceptable (≥ 0.23), medium (≥ 0.49), and high (≥ 0.80)—referred to as the success rate (y-axis). Each method creates 5 structures per AbAg. Success rates are shown for when choosing the highest-confidence structure and the best-by-DockQ structure (Oracle).

### S4: Multiple sequence alignment inputs have little impact on predicted antibody quality

We investigated how MSA input affects the accuracy of antibody structures predicted by Chai-1. Antibody MSAs were generated using MMSeqs2 against the UNIREF30 database under two different conditions. In the first, we used default sensitivity settings with a maximum depth of 300 sequences, referred to as Chai-1 (ShortMSA). In the second, called Chai-1 (DeepTaxMSA), we applied the highest sensitivity setting of 8, increased the maximum MSA depth to 10,000 sequences and using taxonomy IDs to create paired MSAs where possible (for details on taxonomy paired MSAs, see Methods). Compared to predictions made without any MSA input, we observed only minor changes in antibody structure accuracy: a small improvement in median RMSD of 0.05 Å (**Figure S4A**).

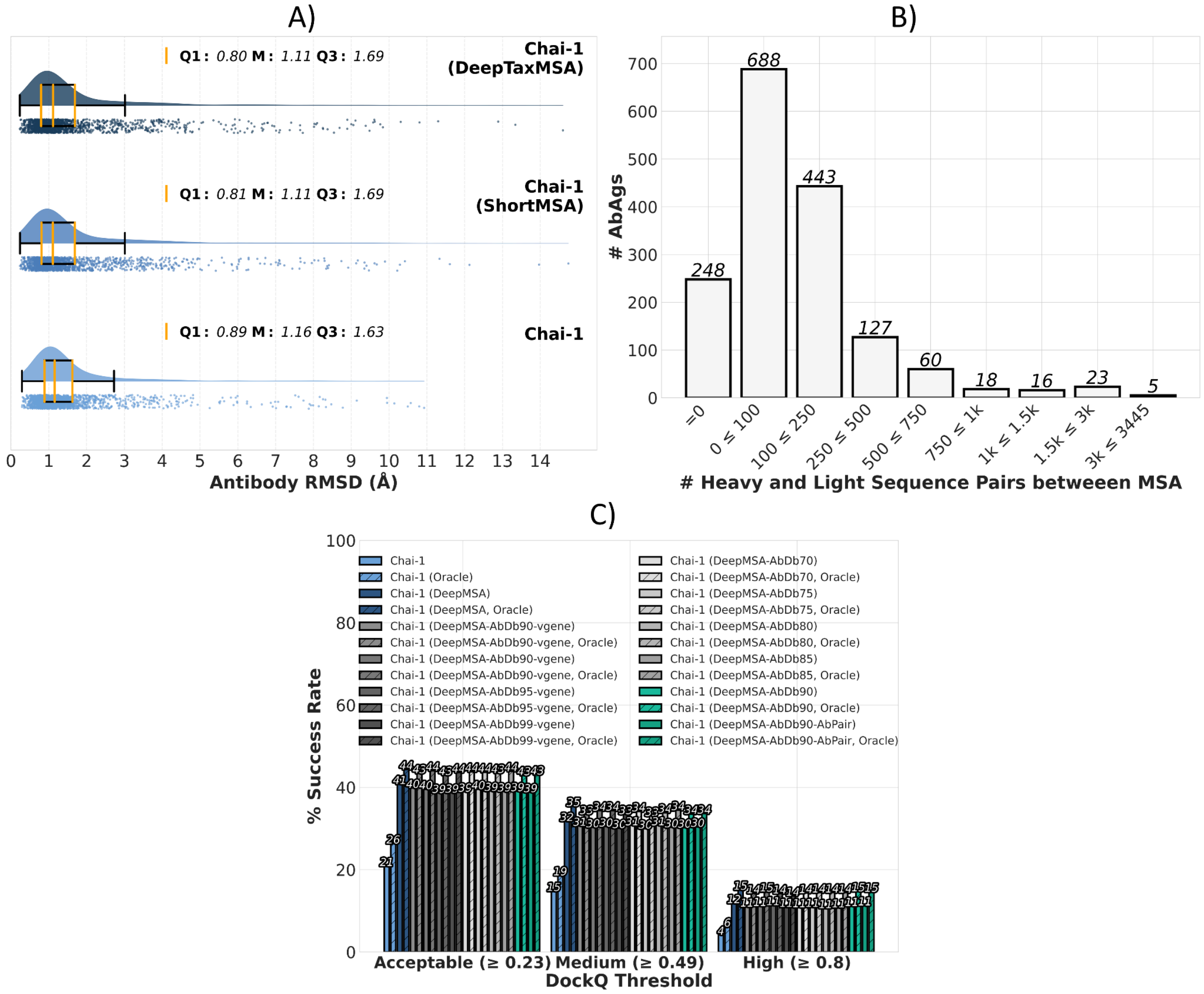


**Fig. S4. Effects of antibody MSAs on predicted antibody and AbAg complex accuracy. A)** Antibody RMSD values (Å, x-axis) are shown for structure predictions across all 1,628 AbAgs, visualized as raincloud plots. For each AbAg, five structures were generated using Chai-1, Chai-1 (ShortMSA), or Chai-1 (MSA), and the RMSD of the antibody chains was computed. Individual predictions are shown as dots; distribution is visualized via density and boxplots. Yellow lines indicate the first quartile (Q1), median (M), and third quartile (Q3). **B)** Histogram showing the number of AbAgs (y-axis) for which light and heavy chain MSAs created from the AbDb90 database contain zero (=0) or a range (x-axis) of sequence pairs originating from the same antibody. **C)** Bar plots showing the percentage of AbAg predictions that meet CAPRI-standard DockQ accuracy thresholds—acceptable (≥ 0.23), medium (≥ 0.49), and high (≥ 0.80)—referred to as the success rate (y-axis). Methods include Chai-1, Chai-1 (MSA), and the 10 Chai-1 (MSA-AbDbX) variants. Each method generates five structures per AbAg; success rates are shown for both the highest-confidence prediction and the best-by-DockQ structure (Oracle).

To investigate whether antibody-specific MSAs could enhance prediction accuracy, we constructed sequence databases composed of paired heavy and light chain sequences from the SABDAB and OAS datasets. These databases allowed for the creation of concatenated MSAs by joining heavy and light chain sequences originating from the same antibody. To assess the practical applicability of this pairing strategy, we analyzed MSAs derived from the AbDb90 database. Among the 1.628 AbAgs, 248 had no matching heavy–light sequence pairs present in the MSA, while the majority (688) contained between 1 and 100 such pairs. This suggests that MSA pairing is possible for a subset of AbAgs, which could impact antibody modelling accuracy (**Figure S4B**).

To systematically evaluate the potential benefit of these antibody-specific MSAs, we generated 10 different paired antibody sequence databases. These were created by varying the MMSeqs2 sequence identity threshold for redundancy reduction (70%, 75%, 80%, 85%, 90%, 95%, and 99%) and by including either full-length antibody sequences or only the regions encoded by V genes. For each of these configurations, antibody MSAs were generated using MMSeqs2 with maximum sensitivity (8) and a depth limit of 10,000. Antigen MSAs were generated from UNIREF30 using the same parameters. These MSA inputs were used for Chai-1 AbAg structure prediction. Across all tested antibody-specific databases, we found no improvement in DockQ-based success rates compared to using UNIREF30 for both antibody and antigen MSAs. We also evaluated whether pairing heavy and light chains within the MSAs (Chai-1 MSA-AbDb90-AbPair) improved accuracy relative to their unpaired counterparts (Chai-1 MSA-AbDb90), but observed no performance gain (**Figure S4C**).

### S5: Re-Evaluating Pocket Restraint Selection Strategies after improving modelled antigen quality with added MSA input

In our main results, we showed that supplying an MSA input to Chai-1 significantly improves the quality of predicted antigen structures (see Figure 4B in Main). Because both DSSP and DiscoTope-3.0 rely on antigen structure, we repeated the restraint-ranking analysis from Supplementary S1 using structure-based scores computed from these improved models.

Specifically, for each of the 1,628 AbAgs in our dataset, we predicted five unbound (no antibody) antigen structures using Chai-1 with DeepTaxMSA and a single seed. DSSP and DiscoTope-3.0 scores were computed using the highest-confidence Chai-1 antigen structure per AbAg.

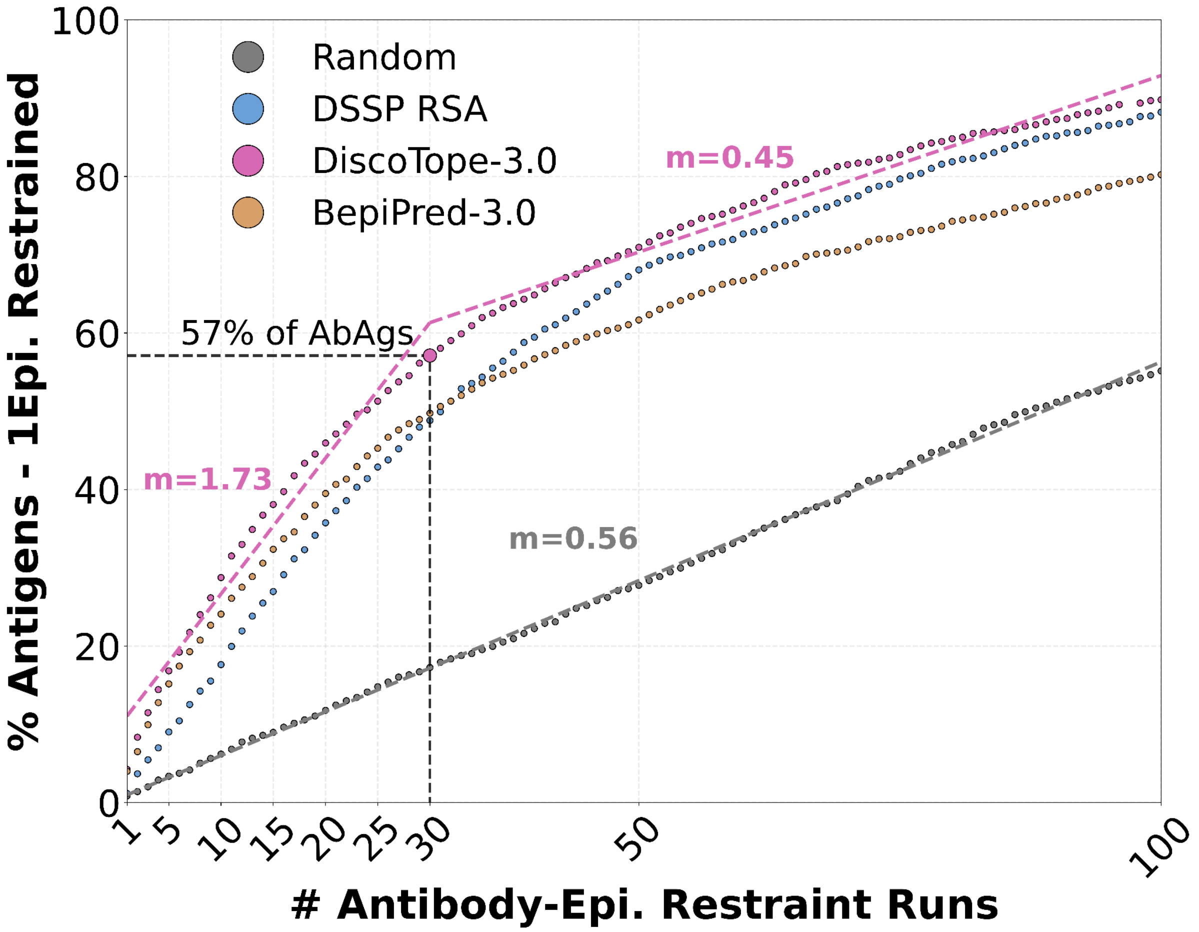


**Fig. S5: Evaluation of epitope restraint selection without AbAg structure prediction.** Percentages of 1,628 AbAg complexes (y-axis) in which the top true epitope contact (i.e., the residue with the most antibody contacts in the solved structure, 1Epi) is included in the restraint, as a function of run number (x-axis). In each run, a new top-ranked residue is selected from a precomputed list, progressively moving down the ranked order. Rankings were derived from BepiPred-3.0 (sequence-based), DiscoTope-3.0, and DSSP solvent accessibility, computed on the highest-confidence Chai-1 (DeepTaxMSA) predicted antigen structure (from five per complex). A random ranking serves as a baseline. Linear slopes (m) are shown for DiscoTope-3.0 (runs 1–30 and 30–100) and Random (runs 1–100).

We found that the improved antigen structures positively impacted the ability of structure-based rankings to prioritize true epitope contacts early. Using the same 30-run milestone from

Supplementary S1, DSSP improved from identifying the top true epitope contact in 43.92% of AbAgs to 48.83%. DiscoTope-3.0 improved more substantially—from 48.16% to 57.12%—surpassing BepiPred-3.0 (49.75%). All methods continued to outperform the random baseline (17.26%).

As before, this benefit appears to plateau after ~30 runs. For DiscoTope-3.0, the rate of discovering new top epitope contacts follows a linear trend with a high slope (m = 1.73) during runs 1–30, but drops to m = 0.45 between runs 30–100, below the random baseline trend (m = 0.56).

### S6: Supplementary figures illustrating BepiPocket and DiscoPocket performance

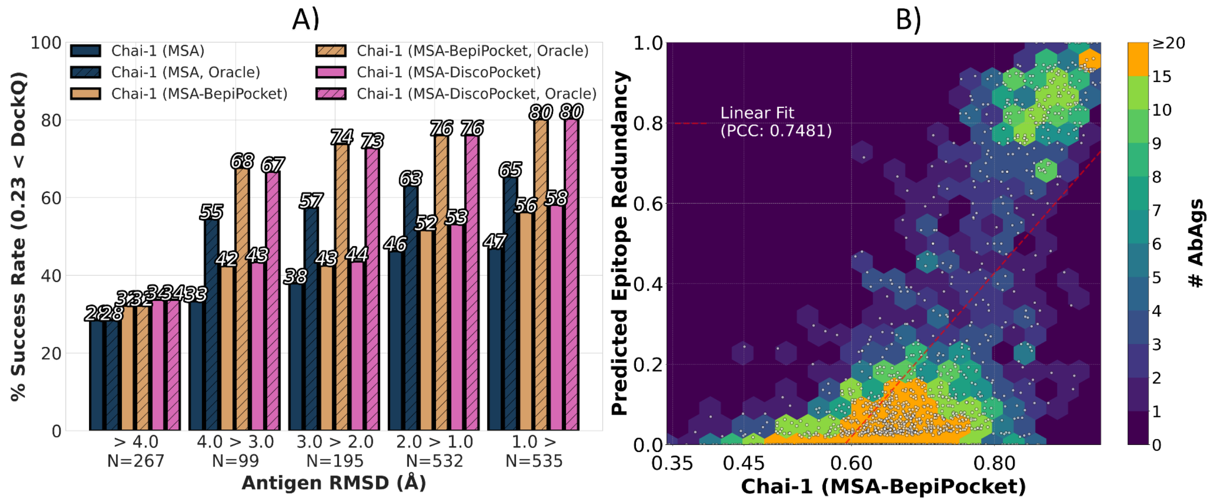


**Fig. S6: Evaluation of the quality Chai-1 structure AbAg predictions for increasingly accurately predicted antigens, and the correspondence between Chai-1 (MSA-BepiPocket) confidence and epitope redundancy for predicted AbAg structures A)** The 1628 AbAgs were grouped by antigen structure quality, measured as the best RMSD (Å) among five Chai-1 predicted antigen structures (x-axis), compared to the solved structure. For each group, we show the percentage of AbAgs with acceptable DockQ accuracy for structures from Chai-1 (MSA), Chai-1 (MSA-BepiPocket) and Chai-1 (MSA-DiscoPocket) Oracle selection. Each method produces 150 structures per AbAg and results are shown for the highest-confidence structure and the best-by-DockQ structure (Oracle).**D)** Median confidence scores of 150 Chai-1 (MSA-BepiPocket) predicted structures per AbAg (x-axis) plotted against corresponding epitope redundancy values (y-axis) for 1,628 AbAgs. Each dot represents one AbAg. To illustrate density, dots are overlaid on 2D hexagonal bins colored by the number of AbAgs per bin (color scale capped at 20).
